# Read trimming is not required for mapping and quantification of RNA-seq reads

**DOI:** 10.1101/833962

**Authors:** Yang Liao, Wei Shi

## Abstract

RNA sequencing (RNA-seq) is currently the standard method for genome-wide gene expression profiling. RNA-seq reads often need to be mapped to a reference genome before read counts can be produced for genes. Read trimming methods have been developed to assist read mapping by removing adapter sequences and low-sequencing-quality bases. It is however unclear what is the impact of read trimming on the quantification of RNA-seq gene expression, an important task in the analysis of RNA-seq data. In this study, we used a benchmark RNA-seq dataset generated in the SEQC project to assess the impact of read trimming on mapping and quantification of RNA-seq reads. We found that adapter sequences can be effectively removed by the read aligner via its ‘soft-clipping’ procedure and many low-sequencing-quality bases, which would be removed by read trimming tools, were rescued by the aligner. Accuracy of gene expression quantification from using untrimmed reads was found to be comparable to or slightly better than that from using trimmed reads, based on expression of *>*900 genes measured by real-time PCR. Total data analysis time was reduced by up to an order of magnitude when read trimming was not performed. Our study suggests that read trimming is a redundant process in the quantification of RNA-seq expression data.

## 1 Introduction

RNA-seq technology is a powerful tool for rapid and comprehensive profiling of expression of genes at a genome scale. The bioinformatic analysis of data generated from this technology however requires significant amount of CPU time and disk storage, because of vast amount of sequence reads generated for even a small RNA-seq experiment. An RNA-seq data analysis includes a number of steps, making it a non-trivial task. There are continuous efforts in the field to try to reduce the complexity of data analysis and improve its efficiency.

Read trimming tools have been developed to remove adapter sequences and bases with low sequencing quality from sequencing reads such as RNA-seq reads, in order to help read aligners to achieve a better read mapping result [1, 2]. Read trimming is the first operation in a sequencing data analysis pipeline that modifies the read sequences produced by a sequencer. The changes it makes to the raw read sequences may impact all the subsequent steps in the analysis pipeline. An important step in analyzing RNA-seq data is the quantification of RNA-seq reads, which assigns reads to genes and counts the number reads assigned to each gene. It is however unclear whether read trimming can improve the accuracy of gene expression quantification. Del Fabbro et al reported that the total number of reads mapping to annotated genes was reduced when read trimming was performed [3]. Didion et al performed a similar study but found that read trimming led to more reads mapping to annotated genes [4]. Williams et al found that read trimming resulted in reduced correlation of RNA-seq data with microarray data [5]. To resolve this issue, more rigorous investigation performed directly on individual genes is required.

Most read aligners have the ability to exclude end bases from the reported alignment of a mapped read if these end bases could not be mapped along with the majority of bases in the read. This process is called ‘soft-clip’ (the excluded read bases are still included in the mapping result but reported as unmapped), which is similar to read trimming but is performed within the read mapping procedure. Soft-clipping is performed solely based on the matching of read bases with reference sequences and it does not require users to provide adapter sequences for adapter removal. However, surprisingly no studies have been carried out to compare soft-clipping to standalone read trimming tools, to the best of our knowledge.

In this study, we first compared read trimming tools to the soft-clipping implemented in the Subread aligner [6, 7]. Then we assessed if mapping and quantification of RNA-seq reads can be improved by read trimming performed prior to mapping. We used a benchmark RNA-seq dataset generated in the SEQC project [8] for this evaluation. The SEQC project also produced real-time PCR data for *>*900 genes, which were used as the truth in our evaluation of quantification accuracy.

## 2 Materials and Methods

A benchmark RNA-seq dataset generated in the SEQC project was used in this study. The SEQC project is the third stage of the MAQC (MicroArray Quality Control) project. Two reference RNA samples were sequenced in the SEQC project: Universal Human Reference RNA (UHRR) and Human Brain Reference RNA (HBRR). This study includes RNA-seq data generated from a UHRR library and a HBRR library. 15 million pairs of 100bp reads was generated from the sequencing of each library.

In the SEQC project, expression levels of *>*1000 genes were validated by the TaqMan RT-PCR technique and 949 of these genes have matched symbols with genes in the RNA-seq data. RT-PCR expression levels of these 949 genes were used as the ‘truth’ of gene expression in the evaluation. The TaqMan RT-PCR data are available from the seqc Bioconductor package [9].

Raw reads or trimmed reads were mapped to the human reference genome GRCh38/hg38 using the ‘align’ function in Rsubread package [7]. ‘align’ is an R wrapper function for the Subread aligner [6]. Soft-clipping is automatically performed by Subread. Reads counts were generated for genes using the featureCounts tool [10]. Rsubread Inbuilt annotation for human genes [7], which is an modified version of NCBI RefSeq gene annotation, was used in the quantification of gene expression. Read counts for each gene were converted to log_2_-RPKM (reads per kilobases per million) expression values and then compared against the RT-PCR gene expression data which is also at log_2_ scale.

When running trimmers to trim reads we tried to keep their default settings where possible. TrimGalore was run with parameters *–illumina -j 8 –paired*. Trimmomatic was run with parameters *PE -threads 8 -phred33 ILLUMINACLIP:TruSeq3-PE.fa:2:30:10 LEADING:3 TRAILING:3 SLIDINGWINDOW:4:15 MINLEN:36* for its ‘adapters and SW’ mode, and run with parameters *PE -threads 8 -phred33 ILLUMINACLIP:TruSeq3-PE.fa:2:30:10 LEADING:3 TRAILING:3 MAXINFO:50:0.5 MINLEN:36* for its ‘adapters and MI’ mode.

All software tools used in this study were executed with 8 CPU threads on a CentOS 6 Linux server with 24 Intel Xeon 2.60 GHz CPU cores and 512GB of memory Versions of these software tools are: Trimmomatic v0.39, TrimGalore v0.6.2 and Rsubread v2.0.0.

## 3 Results

To assess the impact of read trimming on the accuracy of gene expression quantification, we ran two popular read trimmers, Trimmomatic and TrimGalore, on an RNA-seq dataset generated for UHRR and HBRR samples in the SEQC project (See Materials and Methods). Trimmomatic was run with two modes: ‘adapters and SW’ mode and ‘adapters and MI’ mode. Adapter sequences are removed in both modes. In the ‘adapters and SW’ mode, Trimmomatic uses a sliding window approach to remove those read bases that have a low sequencing quality, and in the ‘adapters and MI’ mode a maximum information quality filtering approach is applied. TrimGalore performs adapter removal and quality filtering via calling the Cutadapt tool [2].

We found that 2.3–4.6% of all read bases included in each library were trimmed off, and Trimmomatic removed twice as many bases as TrimGalore (Supplementary Table S1). Total number of successfully mapped bases was reduced by 1.3–4.0% when trimming was applied (Supplementary Table S2). Subread was found to soft-clip 18–29% of bases trimmed off by read trimmers, indicating that a large number of trimmed bases were rescued during read mapping. Out of those commonly removed bases by Subread and a trimmer, 10–27% were found to be adapter sequences and the rest were low-quality bases (Supplementary Table S1). Subread was able to soft-clip almost all adapter sequences (94%) reported and removed by Trimmomatic. TrimGalore reported about six times more adapter sequences than Trimmomatic, but adapter sequences called by TrimGalore are likely to have a high error rate since a lot of them are very short. Nonetheless ~30% of adapter sequences reported by TrimGalore were soft-clipped by Subread (Supplementary Table S3). Put together, Subread was found to be able to effectively remove adapter sequences from the raw reads and rescue a lot of bases with relatively low sequencing qualities which would otherwise be removed by read trimmers. This has led to a nontrivial increase in the number of successfully mapped read bases.

We then examined the impact of read trimming on read mapping results. Read trimming may cause a slight change to the mapping location of a read or cause a read to map to a different exon of the same gene, but this normally would not change the quantification of expression of the gene because the read still overlaps the same gene. We therefore call a read as a concordantly mapped read if read trimming only results in a *<*100bp change in its mapping location or results in the read mapping to an alternative exon from the same gene. We found that *>*98% of reads were concordantly mapped when comparing mapping of TrimGalore trimmed reads and untrimmed reads. Mapping concordance between Trimmomatic trimmed reads and untrimmed reads was ~97%. Mapping concordances between reads trimmed by different trimmers were also found to be ~97% (Supplementary Table S4). The mapping analysis shows that read trimming only affects the mapping of a very small fraction of reads in a library and that the mapping difference between trimmed and untrimmed reads is similar to the mapping difference between reads that were trimmed by different trimmers.

Finally we investigated if read trimming will affect the quantification of gene expression in RNA-seq data. For both trimmed and untrimmed data, we counted the number of mapped reads assigned to each gene using the featureCounts program [10]. Read counts were then converted to log_2_-RPKM expression values for each gene. The SEQC RNA-seq benchmarking study validated the expression of ~ 1000 genes using TaqMan RT-PCR technique [8]. 949 of these genes matched the RefSeq genes and were included in this evaluation. We used RT-PCR expression values of these 949 genes as the truth to assess if read trimming is beneficial to the quantification of gene expression in RNA-seq data. Table 1 shows that performing read trimming before read mapping does not improve the correlation of gene expression values with true values. In fact the correlation has a slight decrease when the reads were trimmed by TrimGalore or Trimmomatic ‘adapters and SW’ mode. This is likely due to the loss of some mappable bases which resulted in less accurate mapping of some reads. Therefore, our evaluation showed that using untrimmed reads to quantify expression levels of genes yielded comparable or slightly better quantification accuracy than using trimmed reads.

**Table 1:** Correlation of trimmed and untrimmed RNA-seq data with the TaqMan RTPCR data. Shown are the coefficients of Pearson correlation between expression levels (log_2_) of 949 genes in UHRR and HBRR samples, measured by the TaqMan RT-PCR technique, and their expression levels (log_2_-RPKM) in RNA-seq data produced by different methods.

| Method | UHRR | HBRR |
| --- | --- | --- |
| No trimming + Subread | 0.851 | 0.870 |
| Trimmomatic-adapters and SW + Subread | 0.850 | 0.870 |
| Trimmomatic-adapters and MI + Subread | 0.850 | 0.871 |
| TrimGalore + Subread | 0.850 | 0.870 |

Performing read trimming also significantly increases the data analysis time (Figure 1). The total running time for producing mapped reads was increased by more than an order of magnitude when using TrimGalore for trimming, compared to no trimming performed. Trimming by Trimmomatic increased the running time by nearly 5 times.

**Figure 1:**
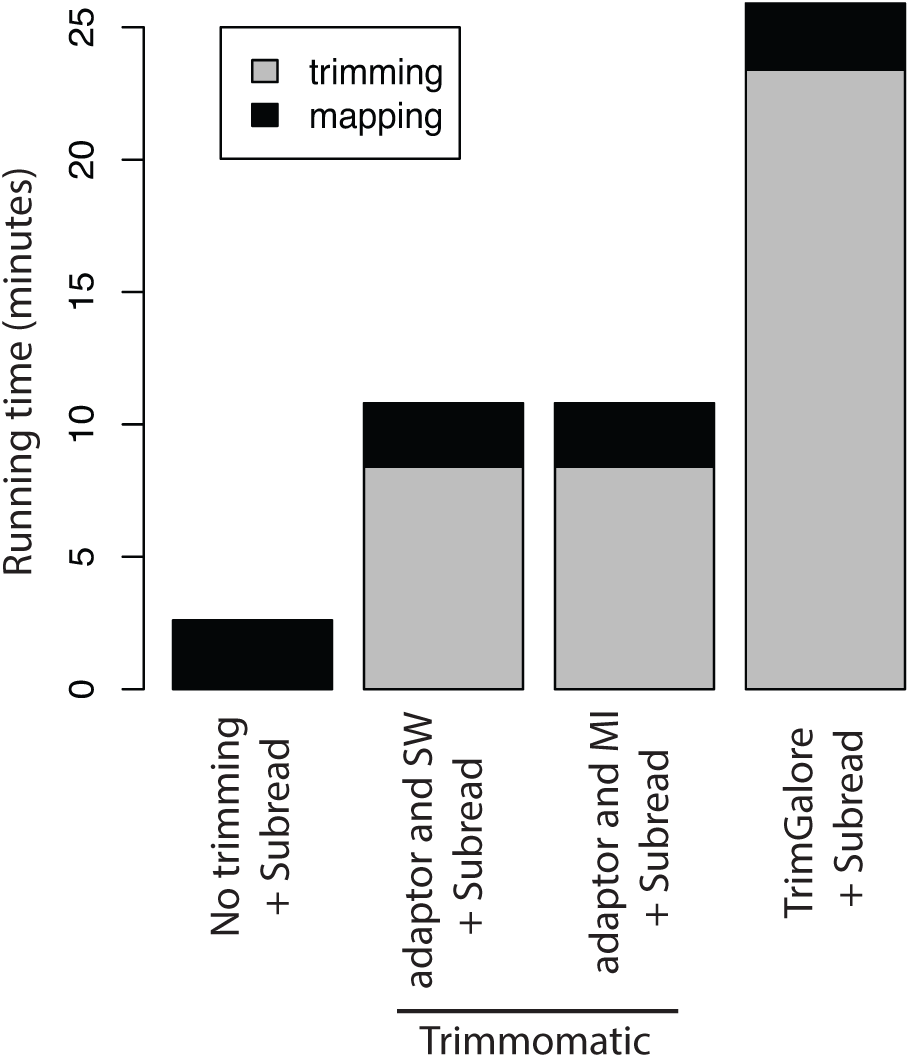
Time cost of different methods running on a UHRR RNA-seq dataset that includes 15 million 100bp read pairs. All software tools were run with 8 CPU threads. Input data to trimming and mapping tools are in gzipped FASTQ format, the standard format of data generated by sequencers.

Furthermore, the amount of disk storage required increased by around 40% due to the need to store trimmed read data (Supplementary Table S5). Read trimming has become a significant computational burden in the analysis of RNA-seq expression data.

## 4 Discussion and conclusion

In this study, we demonstrated that the reference-based soft-clipping implemented in the Subread aligner can effectively remove adapter sequences introduced by sequencers, one of the major goals read trimming tools try to achieve. By leveraging reference sequences, soft-clipping was also found to be able to rescue a lot of low-quality bases. Although we expect that most state-of-the-art RNA-seq aligners are capable of identifying and removing adapter sequences and low-quality bases, Subread has an unique advantage in doing so thanks to its highly flexible and powerful ‘seed-and-vote’ mapping strategy that is more tolerant of adapter and low-quality bases.

We found that read trimming performed prior to Subread mapping did not improve read mapping results and consequently the accuracy of gene expression quantification was not improved either. The quantification accuracy was actually found to be slightly higher when read trimming was not performed. Total RNA-seq quantification time was also found to be reduced by up to an order of magnitude for the datasets used in this study, without read trimming being performed. In conclusion, we recommend that untrimmed reads should be provided for the mapping and quantification of RNA-seq reads.

## 5 Acknowledgements

We thank Gordon K Smyth for suggesting this study.

## 6 Funding

This work has been supported by the Australian National Health and Medical Research Council Project grants 1023454 and 1128609 to W.S.; Walter and Eliza Hall Institute Centenary Fellowship sponsored by CSL to W.S.; Victorian State Government Operational Infrastructure Support and Australian Government NHMRC IRIIS.

## Conflict of interest

none declared.

## References

[1] Bolger,A.M., Lohse,M. and Usadel,B. (2014) Trimmomatic: a flexible trimmer for Illumina sequence data. Bioinformatics, 30, 2114–20.

[2] Martin,M. (2011) Cutadapt removes adapter sequences from high-throughput sequencing reads. EMBnet.journal, 17, 10–12.

[3] Del Fabbro,C., Scalabrin,S., Morgante,M. and Giorgi,F.M. (2013) An extensive evaluation of read trimming effects on Illumina NGS data analysis. PLoS One, 8, e85024.

[4] Didion,J.P., Martin,M. and Collins,F.S. (2017) Atropos: specific, sensitive, and speedy trimming of sequencing reads. PeerJ, 5, e3720.

[5] Williams,C.R., Baccarella,A., Parrish,J.Z. and Kim,C.C. (2016) Trimming of sequence reads alters RNA-Seq gene expression estimates. BMC Bioinformatics, 17, 103.

[6] Liao,Y., Smyth,G.K. and Shi,W. (2013) The Subread aligner: fast, accurate and scalable read mapping by seed-and-vote. Nucleic Acids Res, 41, e108.

[7] Liao,Y., Smyth,G.K. and Shi,W. (2019) The R package Rsubread is easier, faster, cheaper and better for alignment and quantification of RNA sequencing reads. Nucleic Acids Res, 47, e47.

[8] Consortium,S.M.I. (2014) A comprehensive assessment of RNA-seq accuracy, reproducibility and information content by the Sequencing Quality Control Consortium. Nat Biotechnol, 32, 903–14.

[9] Liao,Y. and Shi,W. (2019) seqc: RNA-seq data generated from SEQC (MAQC-III) study. R package version 1.20.0.

[10] Liao,Y., Smyth,G.K. and Shi,W. (2014) featurecounts: an efficient general purpose program for assigning sequence reads to genomic features. Bioinformatics, 30, 923–30.

