## Supplementary Tables S1-S5 for "Read trimming is not required for mapping and quantification of RNA-seq reads"

**Supplementary Table S1.** Comparison of read bases trimmed off by read trimmers and read bases soft-clipped by Subread. Subread was run on the untrimmed reads. Columns 'Trimmed' give the percentages of read bases that were trimmed off by each trimming method, columns 'soft-clipped out of trimmed' the percentages of trimmed bases that were also soft-clipped by Subread and columns 'Adapters in both soft-clipped and trimmed' the percentages of adapter bases found in all the read bases that were commonly removed by a read trimmer and Subread. Adapter percentage of a trimmer was calculated using the adapter bases reported by the same trimmer.

| Method | UHRR | | | HBRR | | |
|---|---|---|---|---|---|---|
| | Trimmed (%) | Soft-clipped out of trimmed (%) | Adapters in both soft-clipped and trimmed (%) | Trimmed (%) | Soft-clipped out of trimmed (%) | Adapters in both soft-clipped and trimmed (%) |
| Trimmomatic-adapters and SW | 4.6 | 19.0 | 10.3 | 4.4 | 17.8 | 11.3 |
| Trimmomatic-adapters and MI | 4.0 | 20.7 | 11.0 | 3.9 | 19.2 | 11.7 |
| TrimGalore | 2.3 | 29.2 | 26.9 | 2.3 | 27.7 | 26.6 |

**Supplementary Table S2.** Percentages of mapped read bases with or without read trimming prior to mapping. Subread was used for mapping of untrimmed or trimmed reads.

| Method | UHRR (%) | HBRR (%) |
|---|---|---|
| No trimming + Subread | 86.4 | 85.5 |
| Trimmomatic-adapters and SW + Subread | 82.4 | 81.7 |
| Trimmomatic-adapters and MI + Subread | 83.2 | 82.3 |
| TrimGalore + Subread | 85.1 | 84.2 |

**Supplementary Table S3.** Total adapter sequences reported and soft-clipped adapters. Columns 'Reported adapters' give the percentage of adapter bases reported by each read trimmer in all the read bases included in a library, and columns 'Adapters that were soft-clipped' the percentage of adapter bases soft-clipped by Subread aligner out of all reported adapter bases.

| Method | UHRR | | HBRR | |
|---|---|---|---|---|
| | Reported adapters (%) | Adapters that were soft-clipped (%) | Reported adapters (%) | Adapters that were soft-clipped (%) |
| Trimmomatic-adapters and SW | 0.10 | 93.8 | 0.09 | 93.5 |
| Trimmomatic-adapters and MI | 0.10 | 93.8 | 0.09 | 93.5 |
| TrimGalore | 0.57 | 32.0 | 0.55 | 29.9 |

**Supplementary Table S4.** Mapping concordance between trimmed and untrimmed reads. The Subread aligner was used in read mapping. Reads were mapped to the human reference genome GRCh38/hg19.

| Method 1 | Method 2 | % reads mapped concordantly | |
|---|---|---|---|
| | | UHRR | HBRR |
| No trimming | TrimGalore | 98.4 | 98.3 |
| No trimming | Trimmomatic-adapters and SW | 96.1 | 96.2 |
| No trimming | Trimmomatic-adapters and MI | 97.4 | 97.5 |
| TrimGalore | Trimmomatic-adapters and SW | 96.2 | 96.3 |
| TrimGalore | Trimmomatic-adapters and MI | 97.8 | 97.8 |
| Trimmomatic-adapters and SW | Trimmomatic-adapters and MI | 97.5 | 97.7 |

**Supplementary Table S5.** Total amount of disk space used by each method. A UHRR library containing 15 million 100bp read pairs was used in this evalution. The consumed disk space includes the storage of raw reads, trimmed reads and mapping results (BAM file). Raw reads and trimmed reads are both in gzipped FASTQ format.

| Method | Disk usage (GB) |
|---|---|
| No trimming + Subread | 5.5 |
| Trimmomatic-adapters and SW + Subread | 7.6 |
| Trimmomatic-adapters and MI + Subread | 7.7 |
| TrimGalore + Subread | 7.8 |
